# Analysis Of Redox Status At The Erythrocyte Level

**DOI:** 10.1101/2022.05.25.493368

**Authors:** Bright Adorbley, Michael Ofosu Preko, Faiza Fodil, Omar Kharoubi

## Abstract

Erythrocytes constitute a significant target for oxidative stress because of their principal function as oxygen-carrying cells. Erythrocytes are constantly exposed to reactive oxygen species as they circulate through the bloodstream. As a result, these erythrocytes have developed a complex antioxidant defense system that comprises both enzymatic and non-enzymatic antioxidants, which mitigates and maintains the erythrocytes in a redox state.

This study aimed to explore the in vitro analysis of the redox status by determining the activities of Catalase, Glutathione S transferase, Thiobarbituric Acid reactive species, and total proteins at the erythrocyte level after exposure to aluminum. The experiment was carried out by incubation of erythrocytes.

The erythrocytes were distributed into sterile Petri dishes in four groups (control group, Al intoxicated group, and intoxicated with Al and treated with quercetin groups (2mg/l and 5mg/l). The experiment was observed over three intervals (24H, 48H, and 72H).

Evaluations were made on oxidative stress and biochemical parameters. Results obtained showed that Al treatment significantly increased the activity of thiobarbituric acid reactive substance and decreased the activity of glutathione S-transferase (GST), activities of catalase, and protein in the erythrocyte of rats. Following administration with quercetin significantly decreased the levels of free radicals and increased the activity of GST, Catalase, and protein activities. Quercetin alleviated the toxic effects of aluminum on the studied parameters.

These results indicate that Al-induced oxidative stress in the erythrocytes of rats. Following administration of quercetin, the activities of the oxidative stress parameters tested were improved significantly, indicating the overall good antioxidant properties of the quercetin. In conclusion, quercetin proved to be an effective antioxidant against free radicals produced in the erythrocytes due to aluminum exposure.

**HIGHLIGHTS:**

1. Erythrocytes are exposed to reactive Oxygen Species as they circulate in the bloodstream.
2. Aluminum induces oxidative stress in erythrocytes of Wistar rats.
3. Aluminum increases the activity of TBARS and decreases the activity of GST and CAT in incubated erythrocytes.
4. Quercetin alleviates the toxic effects of aluminum on biochemical and oxidative stress parameters.

## INTRODUCTION

The balance between pro-oxidants and antioxidants is described as the redox state. Oxidants such as free radicals and other reactive species are constantly produced in the cell. Because it is impossible to eliminate oxidant generation, the cell has evolved many antioxidant systems. Oxidants and antioxidants must be in balance to sustain good health. In the cell, however, maintaining this equilibrium is exceptionally challenging. Oxidative stress is caused when the balance between oxidant and antioxidant is interrupted, shifting the balance toward an oxidized state. The physiopathology of various diseases is linked to oxidative stress. Cardiovascular disease, cancer, diabetes, and various other conditions are among them. (Riera, 2002)

The primary function of red blood cells is gas exchange, wherein they transport oxygen from the lungs to oxygen-poor tissues. The limitation of transporting oxygen is the resulting exposure to high levels of oxidative stress. And therefore, to attenuate oxidative stress-induced damage, the red blood cells have a highly complex antioxidant system that encompasses both enzymatic (catalase and glutathione peroxidase) (Birben et al., 1997) and non-enzymatic factors (glutathione and ascorbic acid)(Ighodaro & Akinloye, 2018). Enzymes in red blood cells and their membranes are ubiquitous. In the cytosol, enzymes such as catalase and glutathione peroxidase minimize the effects of oxidative stress (Rifkind et al., 2003), and a protease complex (the 20S proteasome) degrades damaged proteins. Thus, red blood cells play a crucial role in maintaining oxidative homeostasis, and any dysfunction of that activity is likely to cause detrimental downstream effects.

Aluminum (Al) is one of the most highly distributed metal in the environment, although Al ion has no physiological role in metabolic processes(Exley & House, 2011). Despite being described as ‘redox inactive,’ aluminum is a potent pro-oxidant and may be exerting this activity by forming an aluminum superoxide semi-reduced radical cation, AlO22+. The evidence to support the formation of this complex and its redox activity is burgeoning(Mujika et al., 2011) and suggests that its pro-oxidant activity is significant at concentrations of aluminum which are commonly found throughout the body. Toxic effects of Al arise mainly from its pro-oxidant activity, which results in oxidative stress, free radical attack, and oxidation of cellular proteins and lipids(Exley, 2013).

Quercetin (3,5,7-trihydroxy-2-(3,4-dihydroxy phenyl)-4Hchromen-4-one) has a typical flavonoid structure and contains five hydroxyl groups: it is a dietary flavonoid, which widely exists in plants like a caper, black chokeberry, onion, tomato, and lettuce(Bischoff, 2008). Quercetin has attracted increasing attention due to its anti-obesity, anti-carcinogenic, antiviral, antibacterial, and anti-inflammatory antioxidant properties(Dueñas et al., 2010). Due to its potential health benefits for humans, quercetin has come into the focus of utilization as a nutraceutical ingredient in the food and pharmaceutical industries. The best-described property of quercetin is its ability to act as an antioxidant. It seems to be the powerful flavonoids for protecting the body against reactive oxygen species produced during the normal oxygen metabolism or are induced by exogenous damage(Panche et al., 2016). It has also been shown to be an excellent Vitro antioxidant and a potent scavenger of ROS and RNS(C. G.M. Heijnen et al., 2001).

These antioxidative capacities of quercetin are attributed to the presence of two antioxidant pharmacophores within the molecule that has the optimal configuration for free radical scavenging (Chantal G.M. Heijnen et al., 2002). Moreover, quercetin is suggested to substantially empower the endogenous antioxidant shield due to its contribution to the total plasma antioxidant capacity(Arts et al., 2004).

Therefore, in this context, the objective of our study is to analyze the redox status at the erythrocyte level (in-vitro).

## EXPERIMENTAL PROTOCOLE

### 1.1 Animals and tissue preparation

In this study, Wistar rats weighing (125.83±11.52) were used. The rats were housed under standard conditions with free access to food and water (12hours light/dark, Temperature 22 ± 2°C). The University’s Scientific Committee approved the study protocol.

The rats were sacrificed. Blood was taken by puncture in the abdominal aorta. A quantity of the blood collected was collected into Heparin tubes. Samples taken were then centrifuged at 2000 turns for 10 minutes. The supernatant was removed. We then added a solution of NaCl 0.09% (to wash the blood), after which we centrifuged it again, and the same processes were repeated three times. The supernatant was again removed, and 2ml of NaCl solution was added.

Finally, the pellet was stored in a refrigerator and used to perform biochemical and oxidative stress parameters tests.

### 1.2 Preparation of the aluminum chloride solution (AlCl3)

Aluminum chloride solution was prepared with 20 mg of AlCl3 into 1000ml of distilled water.

#### Preparation of quercetin solution

Two solutions of quercetin solution were prepared:

**Quercetin**1: 2mg of powdered quercetin into 1000ml of distilled water

**Quercetin 2:** 5mg of powdered quercetin into 1000ml of distilled water.

### 1.3 Incubation of erythrocytes

The incubation was carried out in a sterile Hotte (BIOBASE) to avoid any contamination and distort our results; we have made all the necessary equipment available to carry out any difficulties during the experimentation. We set up 36 Petri dishes, divided into four groups (with three repetitions for each group);

**Control Group**: 0.5ml of pellet, 1drop of penicillin, and 10ml of NaCl.

**Al Group**: 0.5ml of pellet, 1drop of penicillin, 10ml of NaCl, 100ul of Al solution

**Al+Q2 group**: 0.5ml of pellet, 1drop of penicillin, 10ml of NaCl, 100ul of Al solution, and 100ul of quercetin solution(2mg/l)

**Al+Q5 group**: 0.5ml pellet, 1drop of penicillin, 10ml of NaCl, 100ul of Al solution, and 100ul of quercetin solution(5mg/l)

They were next placed in an Incubator set to 37°C and CO2 to 5% for a time that ranges from (24, 48H, and 72H).

### 1.4 BIOCHEMICAL ASSAYS FOR OXIDATIVE STRESS

#### 1.4.1 PROTEIN ASSAY

The Lowry Assay Protein(LOWRY et al., 1951) has been the most widely used method to determine the number of proteins in biological samples.

This protein assay was realized based on Lowry et al., and the principle of the technique is based on the biuret reaction but with additional stages and reagents to improve detection sensitivity. Copper reacts with four nitrogen atoms in peptides to generate a cuprous complex in the biuret reaction. Lowry uses the Folin-Ciocalteu reagent, which is phosphomolybdic/phosphotungstic acid. The cuprous ions and the other side chains of tyrosine, tryptophan, and cysteine interact with this reagent to generate a blue-green color that may be detected between 650 and 750 nm. This reaction’s final product is blue in color. The number of proteins in a sample can be measured by comparing the absorbance (at 750 nm) of the Folin reaction’s end product to a standard curve of a selected standard protein solution

#### 1.4.2 THIOBARBITURIC REACTIVE ACID REACTIVE SUBSTANCES (TBARS ASSAY)

The development of malondialdehyde (MDA), which can be detected as Thiobarbituric Acid Reactive Substances, can be caused by oxidizing agents altering lipid structure, resulting in the formation of lipid peroxides (TBARS); the TBARS assay was performed using the technique of Ohkawa et al. The TBARS method, which was first employed in 1978, is still a popular and convenient way to determine the relative lipid peroxide concentration of sample sets such as serum, plasma, urine, and cell lysates and cell culture supernates (Ohkawa et al., 1979).

MDA interacts with TBA in the presence of heat and acid to form a colorful end product (pink chromogen) that absorbs light. At 530-540 nm. The intensity of the resulting color at 532 nm corresponds to the level of lipid peroxidation in the sample. Unknown samples are compared to the standard curve. (Ohkawa et al., 1979).

#### 1.4.3 CATALASE ASSAY

Catalase is present in the peroxisomes of nearly all aerobic cells and protects the cell from the toxic effects of hydrogen peroxide by catalyzing the decomposition of H_2_O_2_ (Deisseroth & Dounce, 1970). In this experiment, the catalase assay was performed using the technique of Sinha (1972). This technique follows the idea that when dichromate in acetic acid is heated in the presence of H_2_O_2_, it reduces to chromic acetate, with perchloric acid as an unstable intermediate. The chromic acetate generated is calorimetrically measured at 610 nm. The presence of dichromate in the test mixture does not affect the colorimetric measurement of chromic acetate since dichromate has no absorbance in this area. For various periods, the catalase preparation is allowed to split H_2_O_2_. The dichromate / acetic mixture injection stops the process at certain time intervals, and the remaining H_2_O_2_ is calculated by measuring chromic acetate calorimetrically after heating the reaction(Sinha, 1972).

#### 1.4.4 GLUTATHIONE-S-TRANSFERASE (GST) ASSAY

Glutathione S Transferase (GST) is an enzyme involved in detoxifying a wide range of compounds and reducing free radical damage in red blood cells(Mannervik & Danielson, 1988). The GST assay in this study was performed according to Habig et al. The principle of GST is that it catalyzes the conjugation of L-glutathione to CDNB through the thiol group of the thiol group glutathione. The reaction product, GS-DNB Conjugate, absorbs at 340 nm. The rate of increase in the absorption is directly proportional to the GST activity in the sample. The reaction is measured by observing the conjugation of 1-chloro, 2,4-dinitrobenzene (CDNB) with reduced glutathione (GSH). This is done by watching an increase in absorbance at 340nm. One unit of enzyme will conjugate 10.0 nmol of CDNB with reduced glutathione per minute at 25°C (Habig et al., 1974).

### 1.5 STATISTICAL ANALYSIS

The results are presented in the form of mean and standard deviation (M+SD), and the analysis of results is analyzed using the student’s t-test (Two samples assuming unequal variances). Values and graphs bearing symbols are significantly different p<0.05.

(*) indicates a significant difference compared to the control group.

(#) indicates a significant difference compared with the intoxicated (Al) group.

## RESULTS & DISCUSSION

**Table 1:** Effects of Aluminium and quercetin on Biochemical and oxidative stress parameters.

| EXPERIMENTAL GROUPS | PROTEINS | CATALASE | TBARS | GST |
| --- | --- | --- | --- | --- |
| **After 24 hours** |  |  |  |  |
| **Control** | 55.49±26.81 | 244.07±139.37 | 820.56±158.23 | 3.58±0.77 |
| **Al** | 41.73±6.82 | 307.04±173.54 | 1905.28±161.41<sup>*</sup> | 2.68±1.34 |
| **Al + Q2** | 59.52±10.45<sup>#</sup> | 149.63±168.58 | 344.24 ±48.57<sup>*#</sup> | 2.68±1.34 |
| **Al + Q5** | 96.03±8.65<sup>#</sup> | 195.56± 73.25 | 744.64±336.68<sup>#</sup> | 4.032±1.34 |
| **After 48hours** |  |  |  |  |
| **Control** | 85.73± 4.92 | 385.37±293.48 | 1023.36±120.88 | 5.78±0.82 |
| **Al** | 67.20 ±4.40<sup>*</sup> | 90.37±62.51 | 1670.24±194.75<sup>*</sup> | 4.66±0.56 |
| **Al + Q2** | 61.41±17.78 | 255±135.51 | 560.56±314.80<sup>*#</sup> | 2.96±0.48<sup>*#</sup> |
| **Al + Q5** | 77.04±17.94 | 403.15±148.93<sup>#</sup> | 625.04±126.13<sup>*#</sup> | 5.15±2.05 |
| **After 72 hours** |  |  |  |  |
| **Control** | 103.49±17.45 | 474.63±126.71 | 1078.48±326.04 | 6.49±0.89 |
| **Al** | 104.77±9.78 | 341.11±50.04 | 1905.28±248.84<sup>*</sup> | 5.062±1.45 |
| **Al + Q2** | 120.45±19.24 | 286.48±117.85 | 1446.64±309.22 | 2.464±0.39<sup>*#</sup> |
| **Al + Q5** | 74.69±9.85<sup>#</sup> | 152.04±101.92<sup>*#</sup> | 749.84±238.68<sup>#</sup> | 2.91±1.39<sup>*</sup> |

Values are represented as mean ± SD for each group. (\*) shows a significant difference of (P<0.05) compared to the control group, and (#) indicates a significant difference of (P<0.05) compared with the intoxicated (Al) group. Student's t-test.

**Figure 1:**
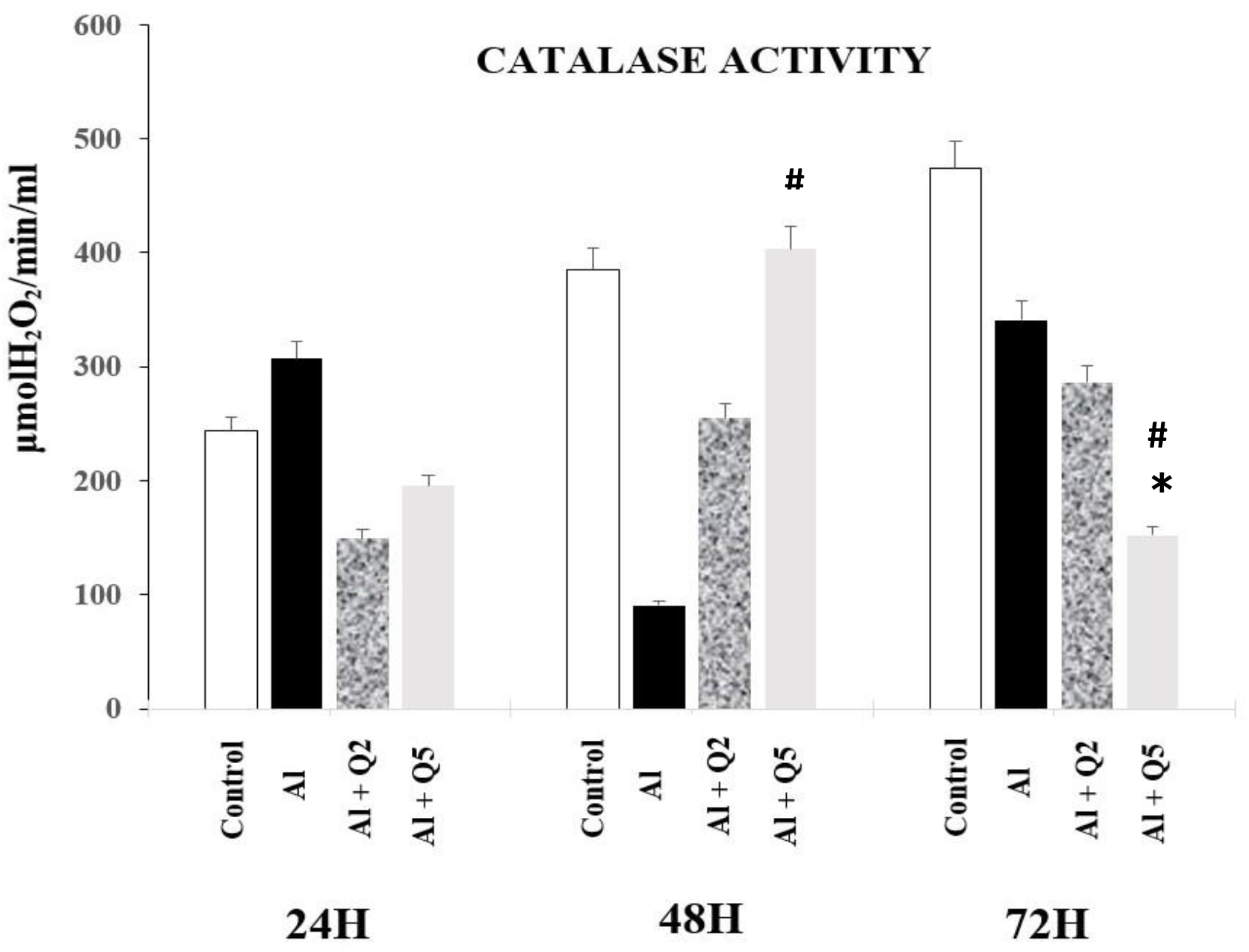
Variation of Catalase activity at the erythrocyte level after Aluminium intoxication and treatment with quercetin. Values are represented as mean ± SD for each group. *:P<0,05 in comparison with the control group; ^#^:P<0,05 compared with the Al group. Catalase activity reflected a significant increase at 24H (25.80%) and a decrease in 48H and 72hH (−76.55% and -28.13%) in the group intoxicated by aluminum compared to the control group. The group intoxicated by aluminum and treated with Q2 highlighted a more significant decrease in catalase activity in all the time variations: 24H,48H, and 72H (−38.69%, -33.83%, and -39.64%, respectively, in comparison to the control group. The group intoxicated with aluminum and treated with Q5 decreased 24H and 72H (−19.89% and -67.97%) in contrast to 48H, which showed an increase of (4.6%) compared to the control group.

**Figure 2:**
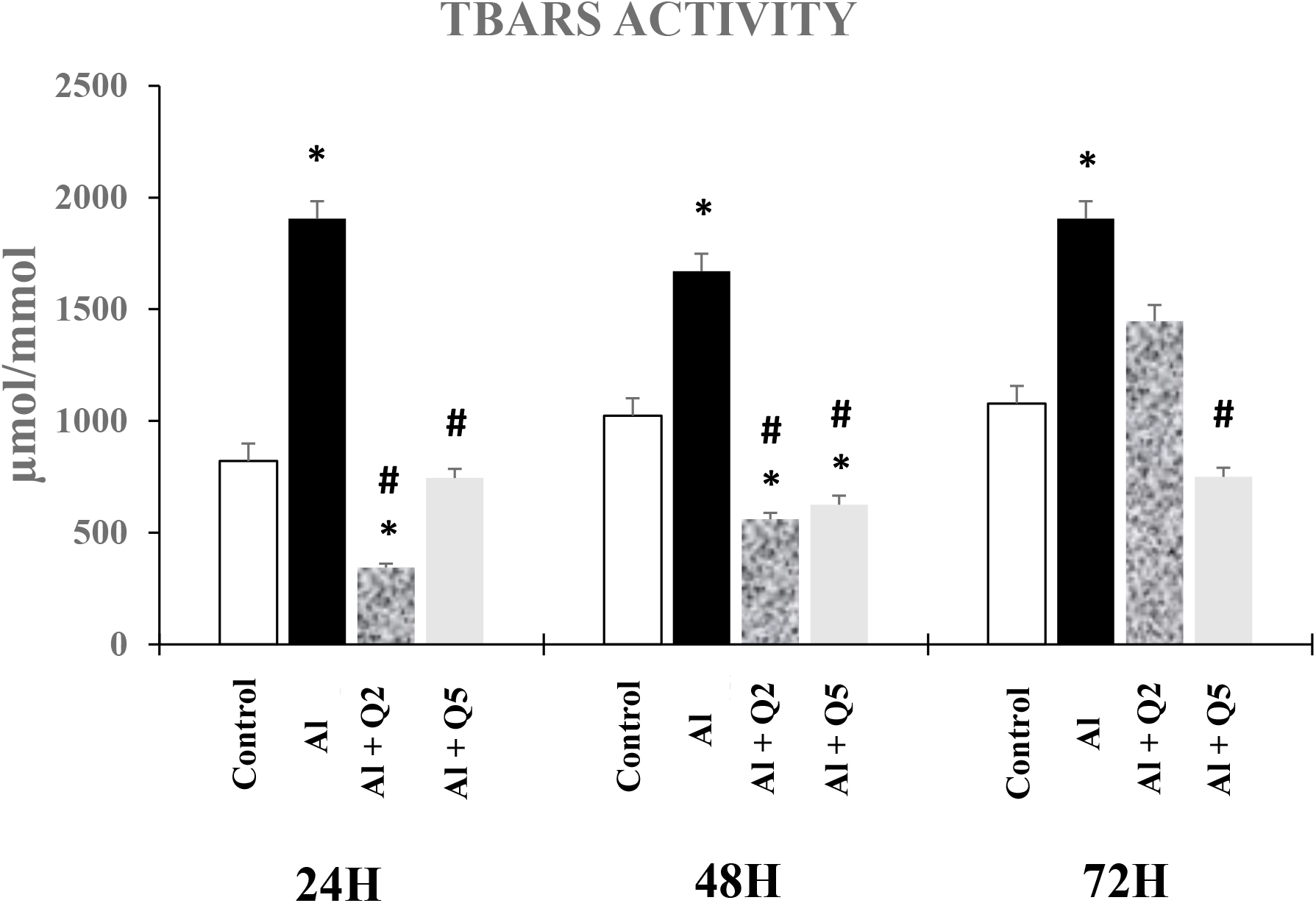
Variation of TBARS activity at the erythrocyte level after Aluminium intoxication and treatment with quercetin. Values are represented as mean ± SD for each group. *:P<0,05 compared to the control group; ^#^:P<0,05 compared with the Al group. TBARS activity revealed a significant increase in all-time variations at 24H, 48H, and 72H (132.19%, 63.21%, and 76.66%) in the group intoxicated with aluminum compared to the control group. The group intoxicated by aluminum and treated with quercetin Q2 displayed a decrease in TBARS activity between 24H and 48H (−58.05% and -45.22%) and, in contrast, a significant increase in 72H (34.14%) compared to the control group. On the other hand, the group intoxicated with aluminum and treated with Q5 quercetin resulted in a significant decrease in all-time various 24H (−9.25%), 48H (−38.92%), and 72H (−30.47%) compared to the control group.

**Figure 3:**
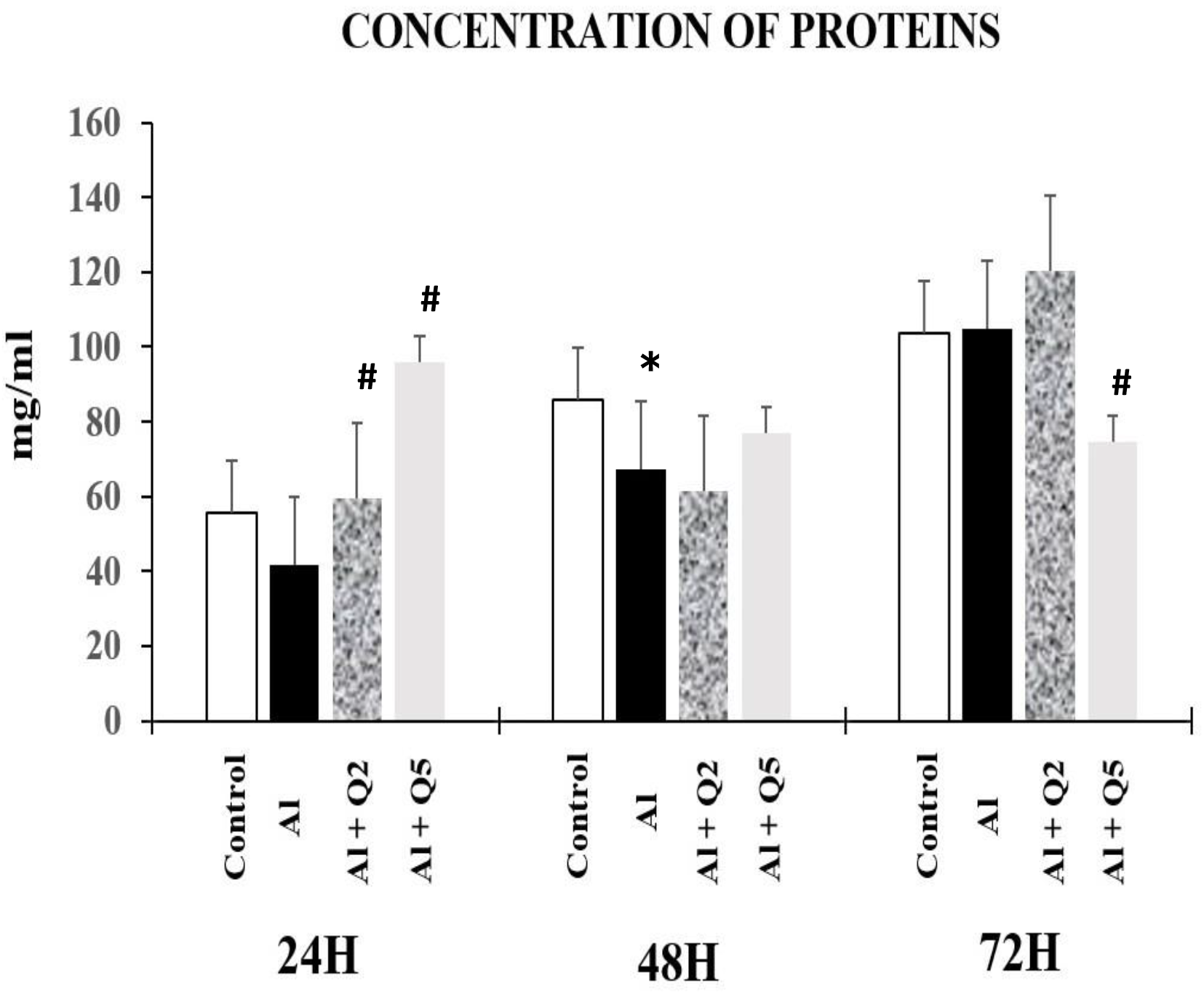
Variation of protein activity at the erythrocyte level after Aluminium intoxication and treatment with quercetin. Values are represented as mean ± SD for each group. *:P<0,05 in comparison with the control group; ^#^:P<0,05 compared with the Al group. Protein content showed a decrease in 24H (−24.80%), 48H (−21.62%), and a slight insignificant increase in 72H (1.23%) in the intoxicated group compared to the control group. The group intoxicated with aluminum and treated with Q2 showed a slight increase in 24H (7.26%) and 72H (16.38%), while there was a decrease at 48H (−28.37%) compared to the control group. Whereas after treatment with a higher concentration of quercetin Q5, we had a highly significant increase in 24H (73.04%) while a decrease in 48H (−10.14%) and 72H (−27.83%) compared to the control group.

**Figure 4:**
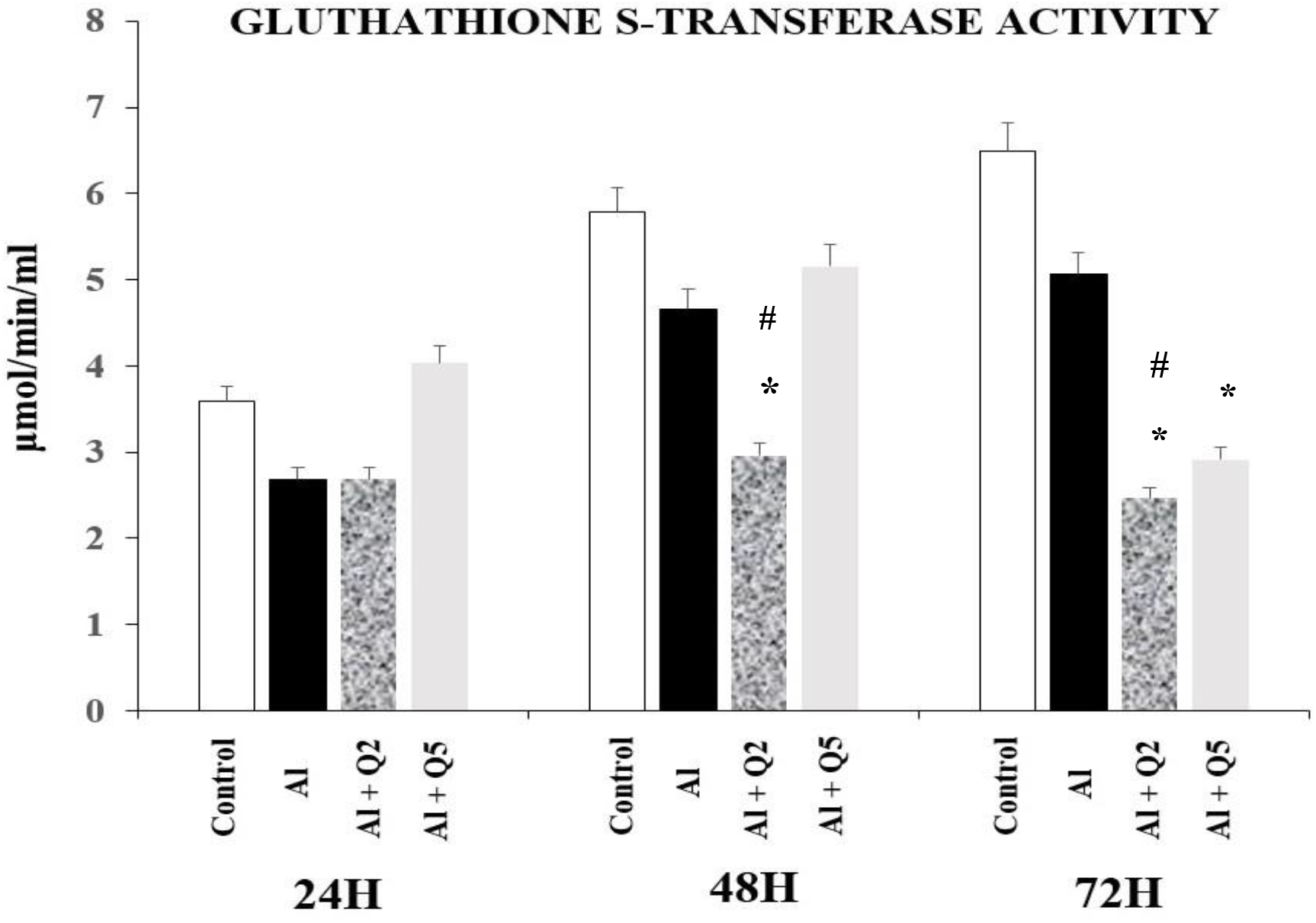
Variation of GST activity at the erythrocyte level after Aluminium intoxication and treatment with quercetin. Values are represented as mean ± SD for each group. *:P<0,05 in comparison with the control group; ^#^:P<0,05 compared with the Al group. GST activity showed a significant decrease in all-time variations 24H (−25%), 48H (−19.38%), and 72H (−22.07%) in the intoxicated group compared to the control group. Compared to the control group, the group intoxicated with aluminum and treated with quercetin Q2 significantly decreased at 24H, 48H, and 72H (−25%, -48.84%, and -62.07%). Whereas after treatment with a higher quercetin Q5, there was a highly significant increase in 24H (12.5%) and contrast to a decrease in 48H and 72H (−10.85% and -55.17%) compared to the control group.

**Table 2:** Effects of Aluminium and quercetin on the number of erythrocytes

| ERYTHROCYTES |  |  |  |  |
| --- | --- | --- | --- | --- |
|  | Control group | Al group | Al+Q2 group | Al+Q5 group |
| **24 hours** | 172100 | 8400 | 194375 | 86920 |
| **48 hours** | 432150 | 52030 | 108000 | 50600 |
| **72 hours** | 404550 | 25525 | 92800 | 3700 |

Values are represented as the mean for each group

**Figure 4:**
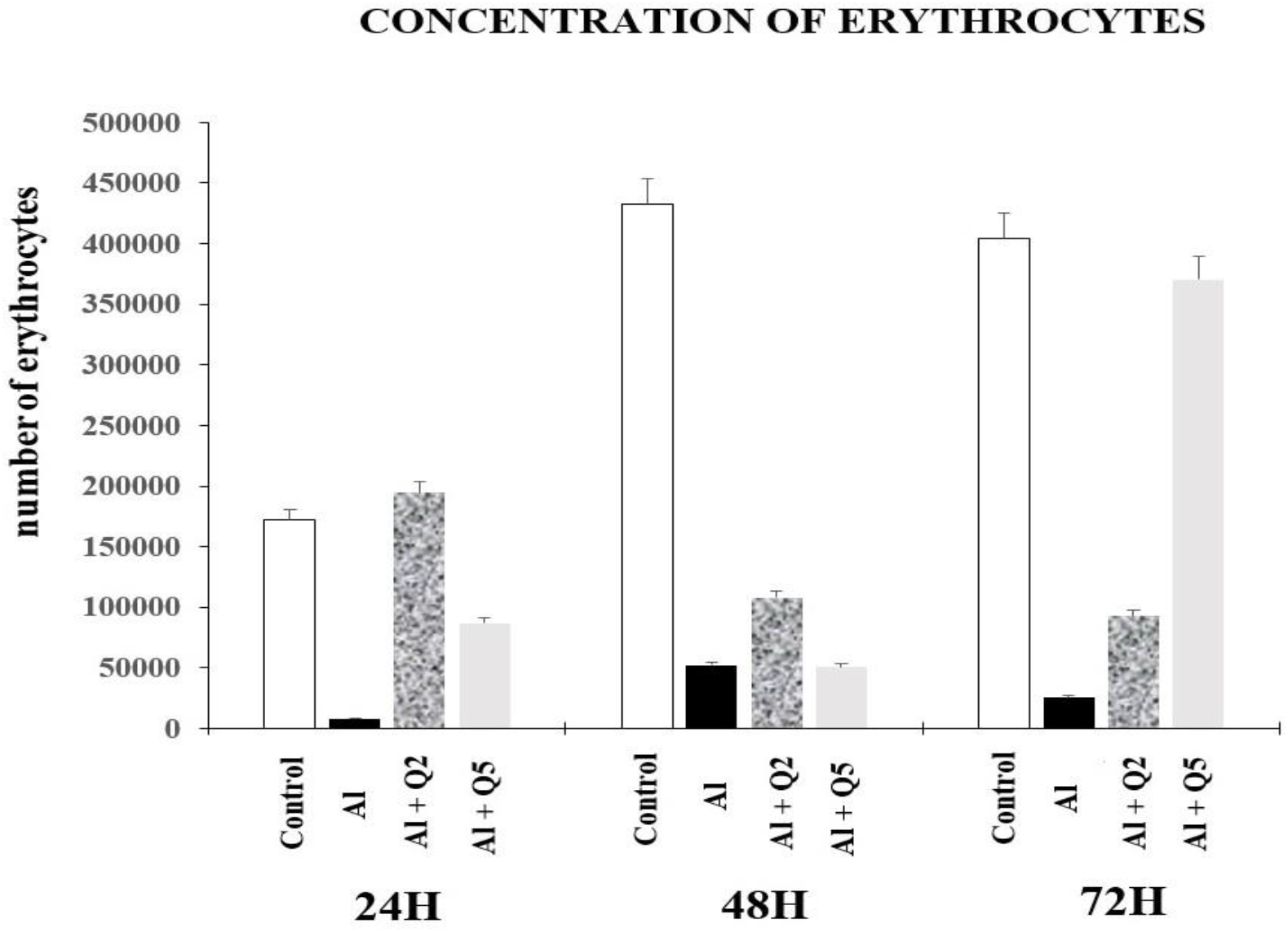
Erythrocytes count after Aluminium intoxication and treatment with quercetin. The concentration of erythrocytes was significantly decreased in all-time variations 24H, 48H, and 72H (−95.12%, -87.96%, and -81.33%) respectively, in the Al-intoxicated, the control group. The improvement was more noticeable in the concentration of erythrocytes in the Al-intoxicated group treated with Q2 quercetin during the 24 hours (12.94%) in contrast to a decrease in 48H (−75.01%) and 72H (−77.06%) compared to the control group. There was again a significant decrease in the number of erythrocytes in all-time variations 24H (−49.49%), 48H (−88.29%), and 72H (−8.39%) in the Al-intoxicated group treated with a higher dose of quercetin(Q5) compared to the control group.

## DISCUSSION

On the effect of intoxication on erythrocytes, we observed a general decrease in the number of erythrocytes across all intoxicated incubations (Aluminium intoxicated-only, and Intoxicated and Quercetin treated Q5mg/L and Q2mg/L) concerning the control incubation. The deleterious effect of Al cannot be restricted merely to the erythroid progenitors since mature erythrocytes might also be affected (Bresnick et al., 2018). This is because Al compounds (chloride and citrate) at concentrations as low as 0.37 μmol/l of Al inhibit erythropoiesis in vitro through a mechanism dependent upon the availability of transferrin to bind to aluminum (Ganz & Nemeth, 2012).

Moreover, electron microscopy analysis of aluminum intoxicated erythrocytes reveals the loss of their typical biconcave shapes and the appearance of abnormal cells such as leptocytes (thin and plain), acanthocytes or schistocytes (spiculated) and target cells, strongly suggesting membrane (Vittori et al., 1999) (Geekiyanage et al., 2019). The alteration of erythrocyte morphology in chronically Al-overloaded rats triggers some questions we asked ourselves (Vittori et al., 2002): Do altered erythroid progenitor cells turn into abnormal erythrocytes? Or, on the other hand, are mature erythrocytes affected by direct contact with Al in the bloodstream during their life span? Our experiment agrees with their work which evaluated the in vitro effects of Al on peripheral erythrocytes by studying their concentration. The results obtained let us assume that Al does affect human erythrocytes in a way that resembles the toxic effects induced in rats by the ingestion of Al (Igbokwe et al., 2020). The incubation of erythrocytes with Al effectively induced changes in their concentration (Vittori et al., 1999). Additional causes of anemia appear to be multi-factorial. They include defective hemoglobin production due to inhibition of the enzymes of heme synthesis, altered erythrocyte membrane structure and fragility, shortening of red blood cell life span due to eryptotic and oncotic injuries, and inadequate iron utilization (Cheng et al., 2018).

Comparisons among the intoxicated incubations allowed us to see a general increase in concentrations of red blood cells in the incubations treated with the Quercetin (Q2 and Q5) concerning aluminum Intoxicated-only incubation.

This could be because of the antioxidant properties of quercetin on the incubated erythrocytes. Quercetin protects erythrocyte membranes against lipid peroxidation (IC50 value = 64±8.7 μM) (Mikstacka et al., 2010). The erythrocyte membrane is prone to lipid peroxidation under oxidative stress that involves the cleavage of polyunsaturated fatty acids at their double bonds, leading to MDA formation (Ayala et al., 2014). This may be a sign of protection to the incubated erythrocytes against hemolysis.

Our work also showed a general decrease in the number of proteins across all intoxicated groups compared to the control group. It has been demonstrated in previous literature that about 2289 unique proteins have been identified in red blood cells (Goodman et al., 2013). However, due to the absence of transcription and translation to replace damaged proteins, erythrocytes have a lifespan of approximately 120 days, after which mature erythrocytes are lost by senescence (Van Wijk & Van Solinge, 2005). An overwhelming amount of hemoglobin in these cells compared to the concentration of all other proteins (Barasa & Slijper, 2014). Early proteomics analyses of erythrocytes failed to provide comprehensive data sets. The presence of hemoglobin and the limitations of proteomics technology to analyze membrane proteins interfered with identifying lower abundant proteins. Progress in the analysis of a cytosolic fraction of erythrocytes was possible using extensive fractionation of proteins (Pasini et al., 2006)(Kakhniashvili et al., 2004) and by reduction of hemoglobin content using peptide libraries (Roux-Dalvai et al., 2008).

ROS can cause protein fragmentation by oxidizing amino acid residue side chains, forming protein-protein cross-linkages, and oxidizing the protein backbone, which could explain a reduction in the number of proteins compared to the control group (Davies, 2016). Under extreme conditions, metal ions profoundly affect cellular protein homeostasis by interfering with their folding process and stimulating the aggregation of nascent or non-native proteins (Hasan et al., 2017). Also, toxic metal ions at the cellular level evoke oxidative stress by generating reactive oxygen species (Li et al., 2016). They promote DNA damage, impair DNA repair mechanisms, impede functional membrane integrity and nutrient homeostasis, and perturb protein function and activity (Tamás et al., 2014). Overall could explain the reduction in the number of proteins in the intoxicated groups compared to the control group.

A significant increase in the 24H and 72H incubations treated with Quercetin Q2mg/L and a highly significant increase in the 24H incubation treated with Quercetin Q5; agrees with works on the antioxidant effects of quercetin on blood (Marunaka et al., 2017). Quercetin shows antioxidant actions via radical scavenging ability and by interacting with antioxidant enzymes, such as heme oxygenase-1 (HO-1), protecting oxidative stress H2O2-induced apoptosis and reducing intracellular ROS production and mitochondria dysfunction (Hahn et al., 2020).

Our work showed a significant increase in catalase concentration in the 24H incubation and a decrease in both 48H and 72H of intoxicated aluminum incubations compared to the control group. Erythrocytes are exposed to oxidative pressure from plasma (mainly hydrogen peroxide – H2O2 and nitric oxide – NO). (Rifkind et al., 2003) Under normal conditions, these cells contain sufficient scavenger enzymes such as Cu, Zn-SOD, CAT, and selenium-dependent GSH-Px to protect themselves from free radical injury. Cu, Zn-SOD catalyzes the dismutation of superoxide (O_2_^٠–^) to H_2_O_2_, which is then independently converted to water by CAT or by GSH-Px (Kurutas, 2016).

So that damage to cells and tissues is avoided, the produced hydrogen peroxide must be immediately converted into other, less-reactive substances. For this purpose, cells often use CAT to quickly catalyze the degradation of hydrogen peroxide into less-reactive oxygen and water molecules (Di Marzo et al., 2018).

Moreover, antioxidant defense enzymes usually act in concert. Thus superoxide dismutase protects catalase and peroxidase against inhibition by O_2_^٠–^, while catalase and peroxidase protect superoxide dismutase against inactivation by hydrogen peroxide (Lei et al., 2015).

This may be concluded as a general decrease in vitro in the activity of catalase over time due to increased concentration of O_2_^٠–^ which gradually inactivates catalase (Bauer, 2015).

However, variations in the results shown by the quercetin treated incubated incubation Q5mg/L (a decrease in 24H and 72H in contrast to 48hours which showed an increase compared to the control group); and Q2mg/L (a more significant decrease in catalase activity in all the time variations: 24H, 48H and 72H in comparison to the control group) showed to some extent the antioxidant of quercetin on the incubated incubations, with a more positive effect in the Q5mg/L incubation concerning the Q2mg/L incubation.

The –OH groups on the side phenyl ring of quercetin are bound to critical amino acid residues at the active site of two enzymes (Ademosun et al., 2016). This way, it has a more substantial inhibitory effect against key enzymes acetylcholinesterase (AChE) and butyrylcholinesterase (BChE), associated with oxidative properties. Our findings also affirm that pre-treatment with quercetin significantly enhances the expression levels of endogenous antioxidant enzymes such as Cu/Zn SOD, Mn-SOD, catalase (CAT), and GSH peroxidase (Hasan et al., 2017).

Glutathione S-Transferases are ubiquitously distributed in nature, being found in organisms as diverse as microbes, insects, plants, fish, birds, and mammals (Gullner et al., 2018). Most of these enzymes can catalyze the conjugation of reduced glutathione (GSH) with compounds that contain an electrophilic center by forming a thioether bond between the Sulphur atom of the GSH and the substrate (Cooper et al., 2020). In addition to conjugation reactions, several GST isoenzymes exhibit other GSH-dependent catalytic activities, including reducing organic hydroperoxides (Allocati et al., 2018). These enzymes also have several non-catalytic functions related to the sequestering of carcinogens, intracellular transport of a broad spectrum of hydrophobic ligands, and modulation of signal transduction pathways (Cho et al., 2001). Many endogenous GST substrates are formed due to the modification of macromolecules by ROS, and the transferases are therefore considered an antioxidant function (Singhal et al., 2015). The GST family, which comprises a relatively high amount of total cytosolic protein, is responsible for the high-capacity metabolic inactivation of electrophilic compounds and toxic substrates (Townsend et al., 2005).

Our work showed a significant decrease in GST concentrations in the intoxicated aluminum incubation in all-time variations compared to the control group. This may be due to the inhibition of GST activity in vitro by the aluminum (Röth et al., 2011), as seen by a significant reduction in concentration across all intoxicated groups (aluminum intoxicated only and treated with Quercetin Q5mg/L and Q2mg/L) concerning the control. Moreover, treatment with quercetin caused a slight increase in concentration compared to the aluminum intoxicated-only group. We observed that the effect was more positive in the Quercetin Q5mg/L treated group compared to the Quercetin Q2mg/L group.

Malondialdehyde (MDA) has been widely used in biomedical research as a marker of lipid peroxidation due to its facile reaction with thiobarbituric acid (TBA). The reaction leads to the formation of MDA-TBA, a conjugate that absorbs in the visible spectrum at 532 nm and produces a red-pink color (Wakimoto et al., 1998). In all the time variations for the intoxicated aluminum group, TBARS activity revealed a significant increase compared to the control group. Lipid peroxidation is a reaction in which free radicals attack lipids, such as ROS and RNS, attack carbon-carbon double bonds in lipids, a process that involves the abstraction of a hydrogen from carbon and insertion of an oxygen molecule; which leads to a mixture of complex products, including, lipid peroxyl radicals, and hydroperoxides as the primary products, as well as malondialdehyde (MDA) and 4-hydroxynonenal as predominant secondary products (Tsikas, 2017).

We observed that the increase in MDA concentration in the aluminum intoxicated group compared to the control group caused a significant rise in concentration. This agrees with abnormal increases in levels of malondialdehyde (MDA) and thiobarbituric acid reactive substances (TBARS) in tissue homogenates of rats exposed to Al (Yu et al., 2019).

Comparing the incubated groups treated with quercetin (Q2Mg/L and Q5mg/L), we observed a significant decrease in concentration in all-time variations of the Q5mg/L group compared to the Aluminium intoxicated group; and conflicting results in the Q2mg/L (a decrease in TBARS activity between 24H and 48H and in contrast a significant increase in 72H of the Q2mg/L compared to the control group). This may prove the ameliorative effect of quercetin in this assay.

Quercetin shows its antioxidant actions, such as protecting cardiovascular cells associated with a decrease in triacylglycerol concentration and an increase in HDL-cholesterol concentration, endothelium-dependent vasodilation due to an increased NO production, and prevention of endothelial cell apoptosis (Pfeuffer et al., 2013). Thus, to some extent, quercetin had a protective effect on in vitro lipid peroxidation (Bustos et al., 2016).

## CONCLUSION

Two Maintaining this status quo in erythrocytes proves to be a delicate task; these cells are constantly overwhelmed with a barrage of free radicals from exogenous and endogenous sources. Red blood cells do well to maintain this balance in vivo by the constant production of more cells to mop out excess free radicals through various defense mechanisms.

Our analysis shows that the maintenance of this balance is limited as, with time, erythrocytes in vitro lose their ground in this battle against free radicals generated by aluminum as there isn’t any production and renewal of red blood cells to combat oxidative stress. Quercetin proved to be a relief to aluminum intoxicated incubated erythrocytes in danger of oxidative stress. Our investigations reveal that improvement in oxidative stress is directly proportional to the concentration of Quercetin treatment added.

## ACKNOWLEDGEMENT

We thank all the team members of the Laboratory of Experimental Biotoxicology, Bio depollution, and phytoremediation of the university of Oran1 Ahmed Ben Bella.

